# Alanine replacements in the structured C-terminal domain of the prion protein reveal conformationally variable regions as major determinants for prion propagation

**DOI:** 10.1101/2023.01.19.524767

**Authors:** Savroop K. Bhamra, Parineeta Arora, Laszlo L. P. Hosszu, Jan Bieschke, Anthony R. Clarke, John Collinge, Parmjit S. Jat

**Author notes:** To whom correspondence should be addressed: Parmjit Jat : MRC Prion Unit at UCL, UCL Institute of Prion Diseases, Courtauld Building, 33 Cleveland Street, London W1W 7FF, UK. Equal contribution (Parineeta Arora, Laszlo L. P. Hosszu). Deceased (Anthony R. Clarke). Joint senior authors (John Collinge, Parmjit S. Jat).

## Abstract

Mutational analysis of the cellular prion protein (PrP^C^) has revealed various regions of the protein that modulate prion propagation. However, most approaches involve deletions, insertions, or replacements in the presence of the wild-type cellular protein, which may mask the true phenotype. Here, site-directed alanine mutagenesis of PrP^C^ was conducted to identify sites particularly a ‘surface patch’ of the protein pertinent to prion propagation in the absence of the wild-type prion protein. Mutations were targeted to the helical, sheet and loop regions of PrP^C^, or a combination thereof and the mutated proteins expressed in PK1 cells in which endogenous PrP^C^ had been silenced. PK1 cells are a clone of mouse neuroblastoma cells that are highly susceptible to Rocky Mountain Laboratory mouse prions. Using the scrapie cell assay, a highly sensitive cell culture-based bioassay for quantifying infectious titres of mouse prions, we found that all mutations within the structured 121-230 domain, irrespective of secondary structure, severely reduced prion propagation. The reduction was most pronounced for mutations within conformationally variable regions of the protein (G123A.L124A.G125A and V188A.T191A.T192A) and those neighbouring or within helix 1 (S134A.R135A.M153A and H139A.G141A.D146A). While mutations G123A and G125A would likely disrupt the structure of the prion fibril, the other mutations are unlikely to cause disruption. Our data therefore suggests that conformationally variable regions within the structured domain of PrP^C^ are the major determinants of prion propagation efficacy.

## Introduction

Prion diseases are fatal transmissible neurodegenerative conditions affecting humans and a range of other mammalian species [1]. Prions consist of a ‘cloud’ of assemblies of misfolded prion protein (PrP) that propagate by recruitment of monomers of the cellular form of the prion protein (PrP^C^) [2-7]. Prion propagation is an autocatalytic process of seeded fibrillisation and fission and involves considerable changes to the overall structure of PrP, from a more α-helical protein to one with a higher β-sheet content [2-5]. Wenborn *et al*, have described a relatively straightforward method for obtaining high-titre, exceptionally pure infectious prions from mammalian brain without Proteinase K (PK) treatment [8]. Recent electron microscopic analyses of exceptionally pure high-titre infectious prions have identified rods comprised of single and twisted pairs of helical protofilament amyloid fibrils where a single PrP monomer is each rung of the protofilament [9-12].

The disease-associated form of PrP, PrP^Sc^, was originally defined in terms of its PK resistance and detergent insolubility, and is pathognomic of prion infection [4, 13]. However, it is now clear that the majority of infectivity in some prion isolates is protease-sensitive [14-16]. While these protease-sensitive infectious forms and cellular PrP^C^ are completely digested upon limited PK treatment, disease-associated forms of infectivity are only partially susceptible to N-terminal digestion by PK and remain detectable by anti-PrP antibodies, such as ICSM18, upon denaturation [17]. Prion infection of a cell can thus be determined by detection of protease-resistant PrP^Sc^ deposits and indeed this forms the basis of the scrapie cell assay (SCA) [18] used to measure prion infectivity in this study.

PrP^C^ has a flexible N-terminal domain (residues 23-122) and a more structured C-terminal globular domain (residues 123-230) comprised of three α-helices (H1, H2 and H3), and a short anti-parallel β-sheet, consisting of two β-strands (S1 and S2) [19, 20]. Helices H2 and H3 are linked by a disulphide bridge. H1 is the shortest of the helices, spanning residues 143-153. It is one of the most immunogenic regions of the protein as numerous anti-prion protein antibodies, including ICSM18, have been mapped to this region and found to recognise PrP^C^ as well as misfolded disease-associated forms upon denaturation [17, 21, 22]. H2 (residues 171-193) and H3 (residues 199-226) are larger in comparison and represent regions of the prion protein where the largest number of pathogenic mutations are clustered [23].

Previous approaches to identify regions within the structured domain of PrP^C^ that influence prion formation have produced conflicting results. All secondary structure components were suggested to influence prion formation via numerous mechanisms. They range from specific protein sequence elements such as the S2-H2-loop [24-27], charge-interactions [28-32], stability [33-37], amyloid-seeding and fibril-forming potential of individual residues [38-41], metal-binding [42], as well as the effect on the protein microenvironment [43-45].

Here, contributions from native amino acids of the cellular prion protein to prion propagation were analysed without interference from either molecular tags or the endogenous prion protein, by targeted alanine replacement, which eliminates sidechains beyond the β-carbon, generally with minimal perturbation of the protein backbone conformation. Alanine replacement has been extensively used to probe the influence of specific amino acid side-chains on bioactivity *in vivo*, and on protein stability, or the folding pathway *in vitro* [46, 47]. It is applicable to a wide range of amino acids, as alanine contains an inert, non-bulky methyl group, and retains the secondary structure preferences of many other amino acids, thus minimally affecting secondary structure. In some cases, alanine replacement may perturb the folding or stability of the modelled protein [48], and care must be taken with regard to the nature of the replaced amino acid. Alanine replacement has previously been used to probe the stability and the folding pathway of PrP^C^ *in vitro* [48, 49], prion replication in cell culture models [50, 51] and also amyloid fibril formation and stability [52-54]. To address the effect of PrP mutations on prion replication *in vivo* without possible interference from wild type PrP, we performed a systematic alanine replacement of spatially proximal residues within the C-terminal domain of PrP (residues 123-231). The mutant proteins were expressed in cells silenced for endogenous PrP expression (PK1-KD cells) [55, 56], in contrast to previous studies which have used cells expressing endogenous PrP, or chronically infected N2a (ScN2a) cells [50, 57]. The aim was to determine if there was a critical ‘surface patch’ of the protein or whether the hydrophobic core of PrP dominated efficient prion propagation.

## Results

### Alanine mutagenesis of the mouse prion protein region 123-230

Mutations to alanine were introduced at various sites in mouse PrP (PDB entry 2L39) [58] to identify a ‘surface patch’ required for efficient prion propagation. Nineteen constructs were generated: sixteen targeted surface amino acids within the globular domain, and three targeted core hydrophobic residues (Fig. 1A). Mutations to surface residues were conducted in triplicate at spatially proximal sites in the 3D structure to create a mutated ‘surface patch’ (Fig. 1B, shown in yellow). Within the core hydrophobic regions of the protein, replacements to alanine, as well as introduction of polar residues, were undertaken at specific sites aimed at reducing protein stability [59] and promoting protein unfolding [60] (Fig. 1B, shown in blue).

**Figure 1.**
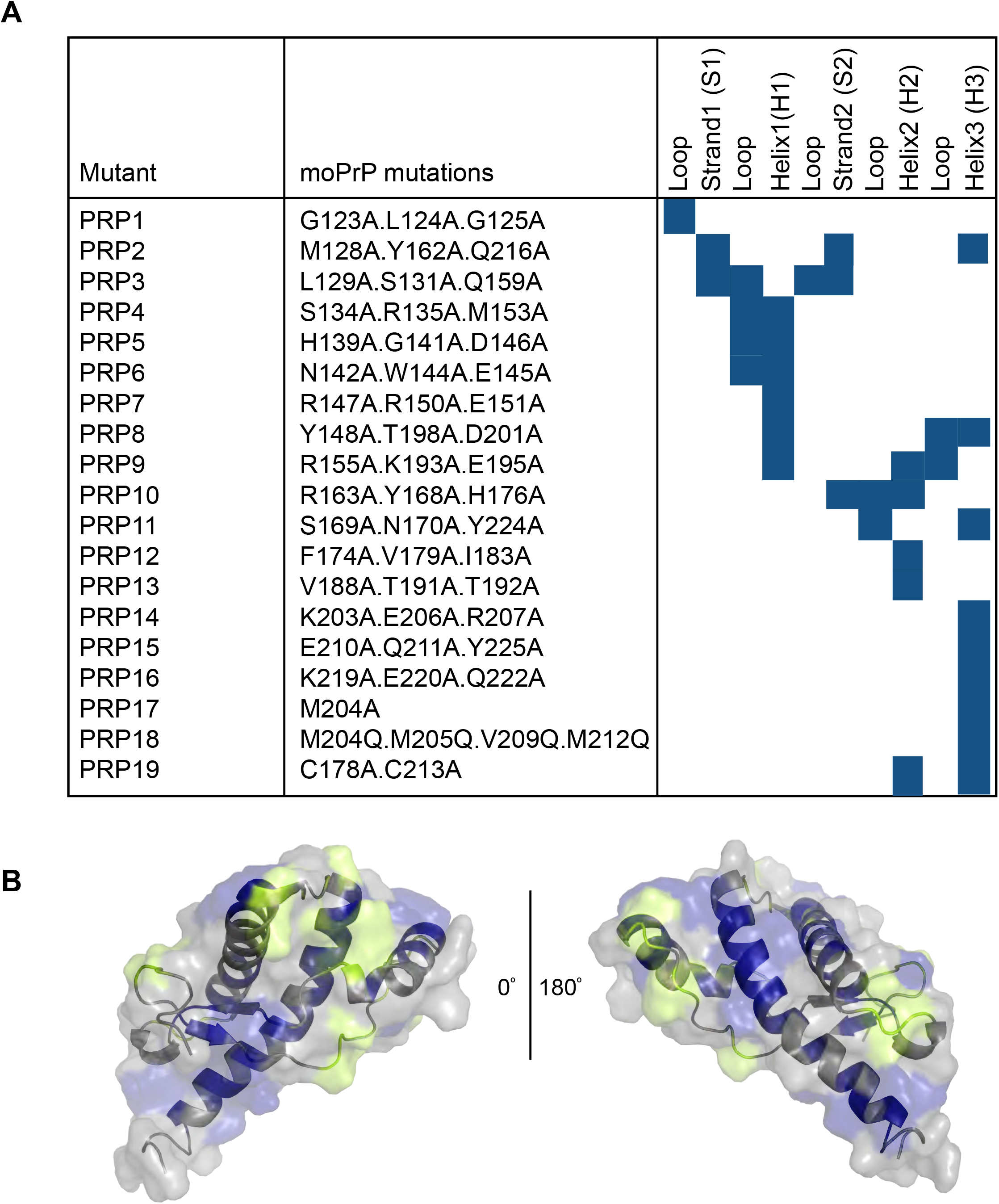
Mutagenesis of the mouse prion protein structured region (123-230) MoPrP mutations were created in the structured region of the protein, largely as triple alanine replacements on surface residues. A. The full list of mutations assayed is detailed in a tabular format, listed by mutation (N-to C-terminal), nomenclature used (PRP1-19) and secondary structure elements affected (loop/strand/helix). B. Targeted mutations mapped on to PDB entry 2L39; images created in PyMol software (PyMol Molecular Graphics System, Version 2.0, Schrödinger LLC). Residues that remained unchanged are shown in grey; mutated surface residues are shown in yellow and mutated buried residues are shown in blue.

The mutations were stably expressed by retroviral infection of PK1-KD cells in which endogenous prion protein expression has been stably silenced [55]. As the level of PrP^C^ is reduced by about 90% in these cells, they are resistant to prion infection, but upon reconstitution with wild-type mouse PrP (moPrP^WT^) regain full susceptibility to prion infection, as determined by SCA [18]. We reconstituted PK1-KD cells with the nineteen mutants (PRP1-19), derived bulk cultures of stably transduced cells, analysed them for PrP expression.

### Mutant PrPs were expressed and trafficked to the cell surface

In all the mutants, PrP was expressed and localized to the cell surface (Fig. 2A). The pattern of expression was the same as for PrP^C^ in PK1 cells, a derivative of N2a cells, that have previously been shown biochemically to express PrP on the cell surface [61]. Although the expression level was variable, all mutants except PRP8, 12, 13 and 19 were expressed at levels equivalent to endogenous PrP^C^ and exhibited the same pattern of three bands corresponding to endogenous PrP^C^ that is non-, mono- and di-glycosylated PrP in the parental PK1 cells (Fig. 2B). Mutants PRP8, 12, 13 and 19 did not express di-glycosylated PrP but predominantly expressed mono-glycosylated PrP. The mono-glycosylated PrP from PRP9, 12, 13 and 19 migrated differently from the mono-glycosylated PrP from wild type and other mutant PrPs (Fig. 2B). Nevertheless, the mutant PrPs were all expressed on the cell surface (Fig. 2A) indicating that the lack of di-glycosylation and aberrant migration did not prevent them from being trafficked to the cell surface.

**Figure 2.**
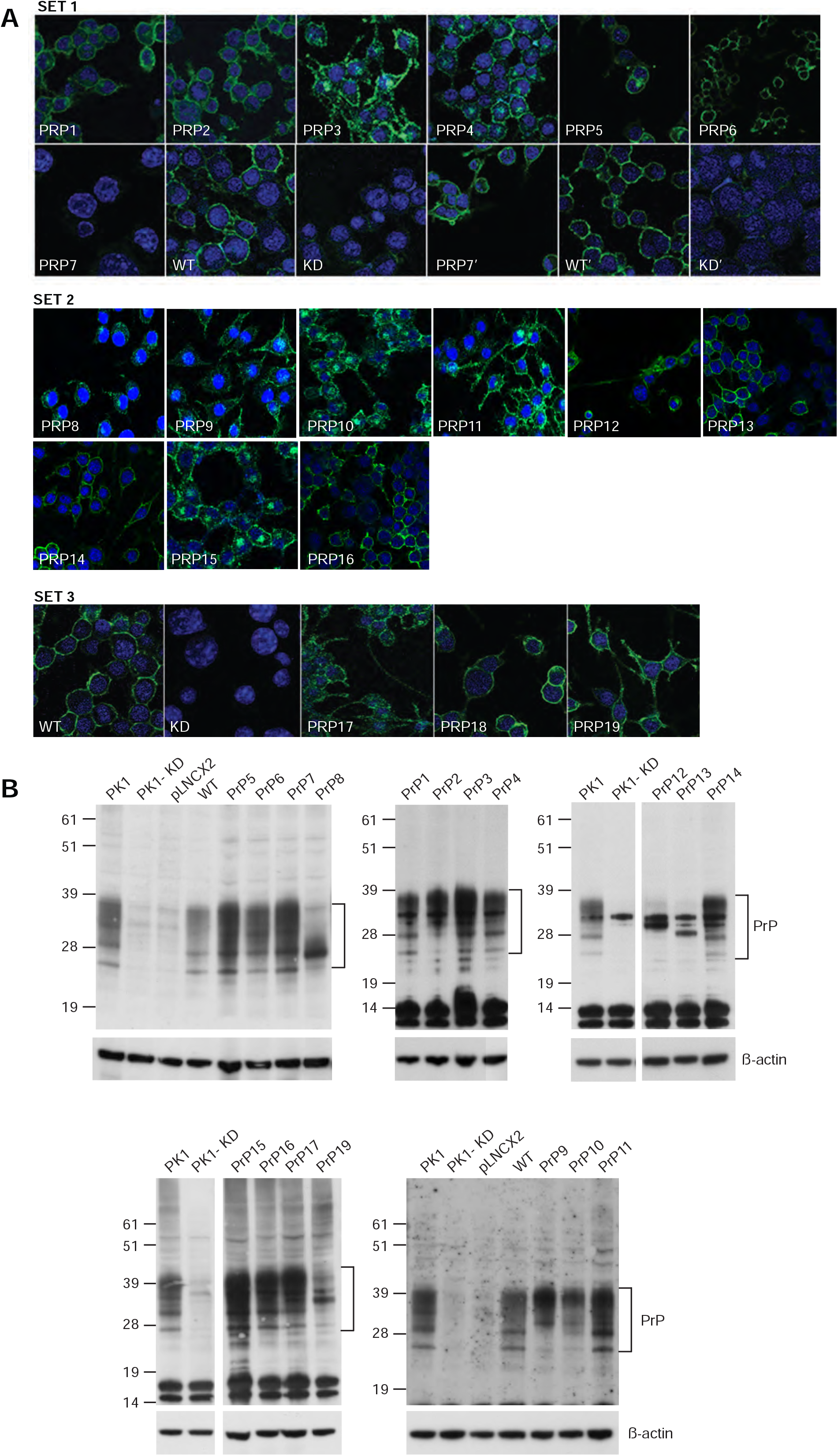
Analysis of PrP expression. A. Immunofluorescence analysis of PK-KD cells reconstituted with the wild-type protein [moPrP^WT^ (WT)] or alanine mutants. SET1: analysis of PRP1-7 using anti-PrP antibody ICSM18 and ICSM35 for PRP7, PK1-KD and KD cells (indicated with ‘) ; PRP7 was recognized by ICSM35 but not detected by ICSM18 (antibody epitope within mutated region). SET2: analysis of PRP8-16 using anti-PrP antibody ICSM18. SET3: analysis of PRP17-19 using anti-PrP antibody ICSM18. All reconstituted KD lines expressed moPrP on the cell surface. DAPI nuclear stain; moPrP, green. Magnification at x40. B. Western blot analysis of lysates prepared from bulk cultures of PK1-KD cells stably reconstituted with the indicated mutants. PK1 correspond to lysates prepared from PK1 cells, KD from PK1-KD cells, pLNCX2 from cells reconstituted with the empty pLNCX2 retroviral vector and WT from cells reconstituted with wild type mouse PrP. Immunoblot analysis was carried out using the anti-PrP antibody ICSM35. The data presented are representative of multiple western blots where a large number of lysates were analysed and show that reconstituted cells express varying levels of PrP isoforms.

Mouse PrP is glycosylated at asparagine180 and196. Even though neither of these residues was mutated in our study, the T198A mutation in PRP8 is part of the consensus sequence for N-linked glycosylation of asparagine 196 and therefore likely to affect its glycosylation. PRP8 did not express di-glycosylated PrP but the mutant PrP was still trafficked to the cell surface (Fig. 2A) in accordance with a previous study that showed that a T198A mutation in PrP eliminated glycosylation of this site but did not alter its trafficking to the cell surface [62]. Mutants PRP12, 13 and 19 also have altered glycosylation and aberrant migration but were trafficked to the cell surface. PRP12 contains V179A.I183A whereas PRP19 contains C178A, very close to asparagine 180; it is possible that these mutations perturb glycosylation at this residue. PRP9 contains K193A.E195A that are adjacent to asparagine 196, but glycosylation was unaffected. However, the monoglycosylated PrP migrates aberrantly in comparison to mono-glycosylated wild type PrP.

Some of the mutants (PRP 3, 4, 10, 11 and 15) affected PrP trafficking leading to partial intracellular accumulation as observed previously [63, 64]. It has been well documented that GPI-anchored cell surface expression of PrP^C^ is necessary for replicating infectivity (62), but the actual level of PrP^C^ expression is not important. Enari *et al* found that N2a/Bos2 cells, a clone of susceptible cells, expressed PrP^C^ at the same low level as the original N2a cells whereas a resistant cell line expressed it at a 10 times higher level [65]. PrP^C^ over-expression did not increase susceptibility either leading them to conclude that it is a prerequisite for prion propagation but other factors are essential for susceptibility to prion infection (64). Marbiah *et al* also showed that overexpression of PrP^C^ did not render revertant cells susceptible to prion infection and did not increase the susceptibility of prion permissive cells [66].

### Mutations to PrP^C^ at surface-exposed sites severely inhibit prion propagation

PrP^C^ mutations were analysed in three sets across residues 123-225. Set 1 (PRP1-7: residues 123-151; Fig. 3) and 2 (PRP8-16: residues 148-225; Fig. 4) primarily covered surface residues, whereas set 3 (PRP17-19; Fig. 5) targeted buried hydrophobic residues. Propagation of RML prions was measured as the number of prion-infected cells detected per 25,000 PK1-KD cells reconstituted with each construct. Cells were passaged through six rounds by splitting biweekly 1:8. The number of infected cells in splits 4-6 was defined as the “spot number”, and determined by counting the number of PrP^Sc^-positive (indicating prion-infected) cells identified, using anti-PrP ICSM18 antibody.

**Figure 3.**
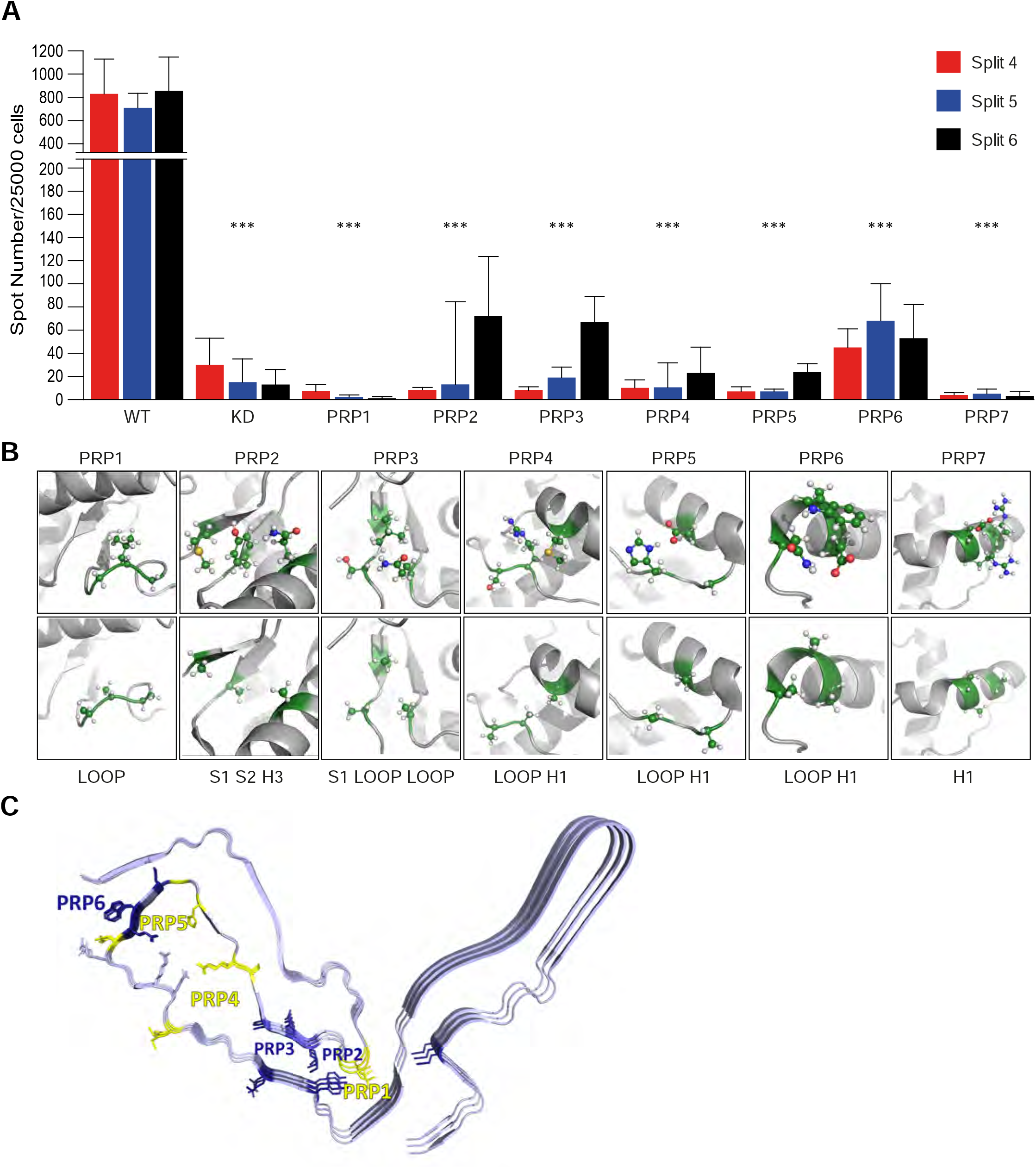
RML prion infection of PK1-KD cells expressing mutations in region 123-151 (PRP 1-7) A. PK1-KD cells reconstituted with either the wild-type protein or alanine mutants of moPrP were challenged with RML prions and assayed for their ability to propagate prions via SCA for three consecutive passages. *** p≤0.0001 calculated in a one-way ANOVA with a Bonferroni correction for multiple comparisons. B. Mutations in PRP1-7 are depicted on PrP^C^ (PDB entry 2L39), with target residues for mutation highlighted. C. Mutations in PRP1-7 are depicted on the RML prion structure (PDB entry 7QIG), with target residues color-coded for relative inhibition of prion replication. PRP7 is uncoloured, as it affects the epitope of the ICSM18 antibody and so could not be assayed for its effect on prion propagation in SCA. All images were created in PyMol.

**Figure 4.**
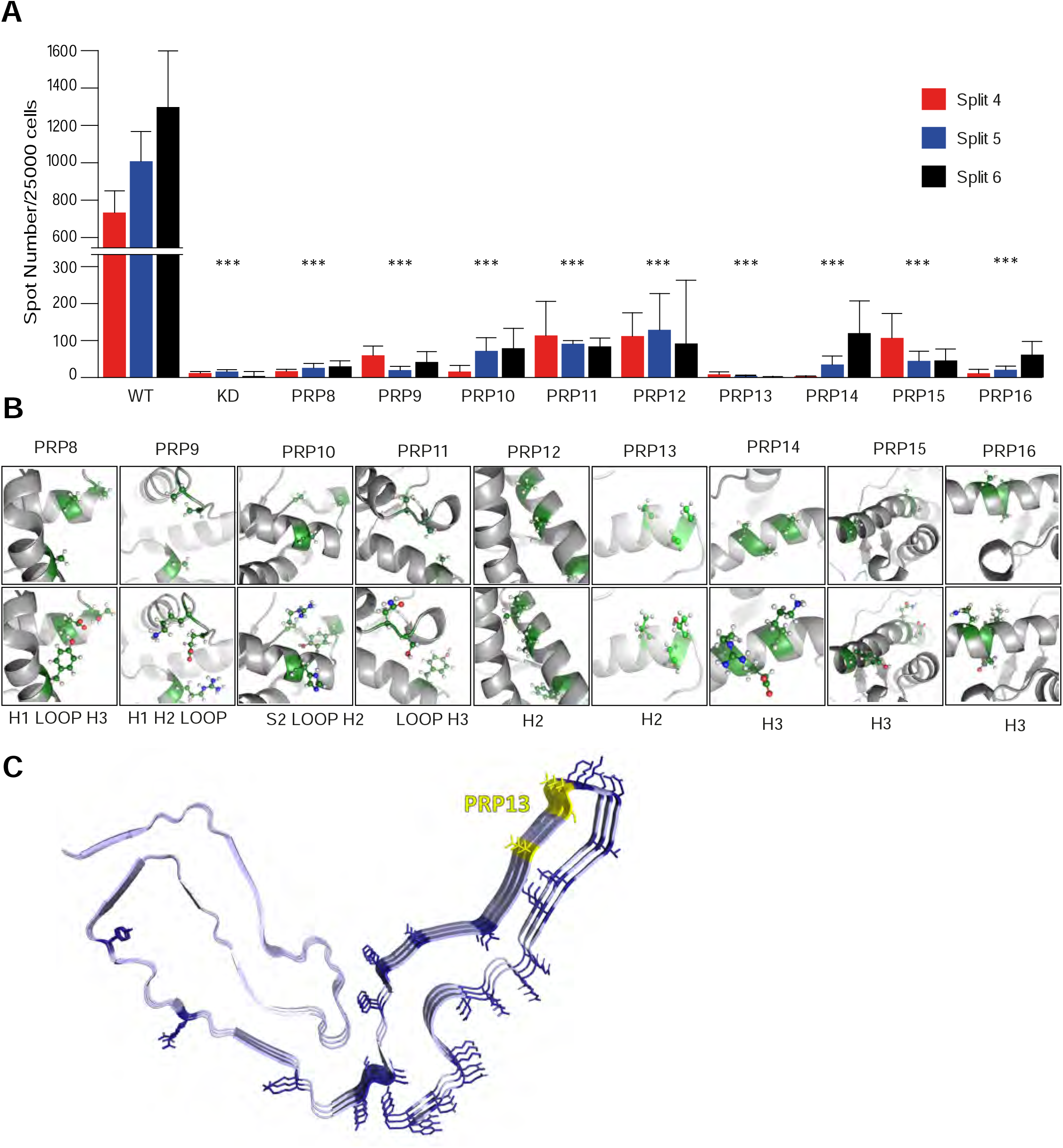
Infection of PK1-KD cells expressing mutations in region 148-222 (PRP 8-16) with RML. A. PK1-KD cells reconstituted with either the wild-type (WT) protein or alanine mutant variants of moPrP were challenged with RML prions and assayed for their ability to propagate prions via SCA for three consecutive passages. *** p≤0.0001 calculated in a one-way ANOVA with a Bonferroni correction for multiple comparisons. B. Mutations in PRP 8-16 are depicted on PrP^C^ (PDB entry 2L39), with target residues for mutation highlighted. C. Mutations in PRP 8-16 are depicted on the RML prion structure (PDB entry 7QIG), with target residues color-coded for relative inhibition of prion replication.

**Figure 5.**
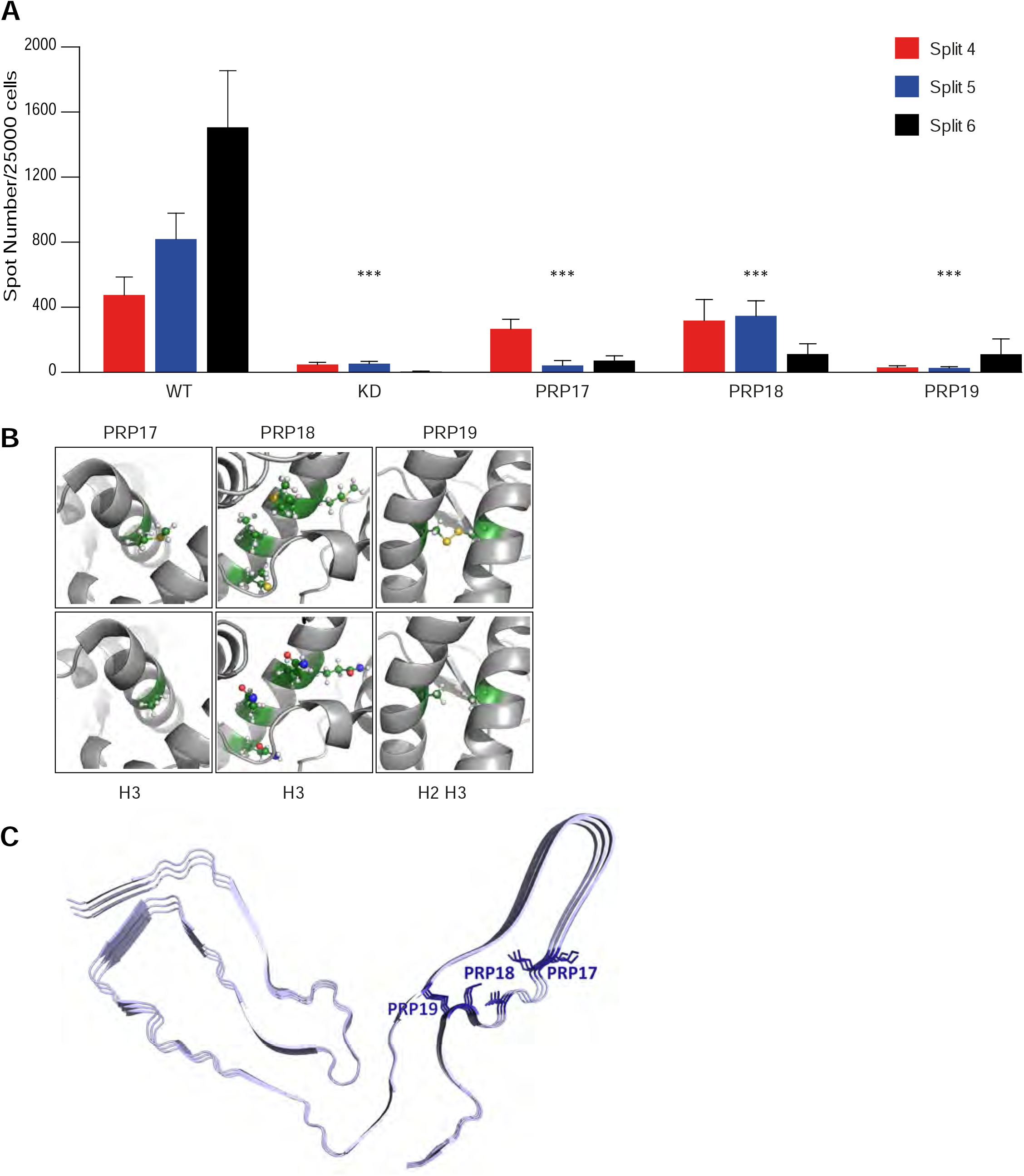
RML prion infection of PK1-KD cells expressing mutations in region 204-213 (PRP 17-19) A. PK1-KD cells reconstituted with either the wild-type protein or mutant variants of moPrP were challenged with RML prions and assayed for their ability to propagate prions via SCA for three consecutive passages. *** p≤0.0001 calculated in a one-way ANOVA with a Bonferroni correction for multiple comparisons. B. Mutations in PRP17-19 are depicted on PrP^C^ (PDB entry 2L39), with target residues for mutation highlighted. C. Mutations in PRP17-19 are depicted on the RML prion structure (PDB entry 7QIG), with target residues color-coded for relative inhibition of prion replication.

Overall, fewer prion-infected cells were observed in cells expressing moPrP mutations in set 1 (PRP1-7, region 123-151), compared to set 2 (PRP8-16, region 148-225); both sets however, comprised specific mutations that permitted only very low-level propagation or completely abrogated propagation (Figs. 3A and 4A). For any mutation in set 1 and 2, the level of prion propagation was a fraction of that for moPrP^WT^ (800-1000 spots), suggesting that regions 123-151 and 148-225 are highly sensitive to mutations at surface sites, as substitution of up to three residues within the native sequence to alanine resulted in a ten-fold decrease in propagation. Furthermore, mutations to conformationally variable regions reduced propagation to background levels in PRP1 (Fig. 3A) and PRP13 (Fig. 4A). PRP7 also produced very few spots, but this is due to this mutant protein not being recognised by the anti-PrP ICSM18 antibody (Fig. 2A).

### Mutants PRP1 and PRP13 abrogate prion propagation

The lowest propagation was observed for those mutants that exclusively targeted conformationally variable regions: PRP1 (G123A.L124A.G125A), N-terminal to S1 (Fig. 1A) and PRP13 (V188A.T191A.T192A), at the C-terminus of H2 (Fig. 1A). Prion propagation in PK1-KD cells reconstituted with these two mutants was akin to background PK1-KD levels if not lower by the last split, indicating complete abrogation of prion propagation (Fig. 3A and 4A). The residues mutated in PRP1 (123-125) display a high degree of conformational flexibility, as shown by NMR relaxation measurements [67, 68], as they immediately proceed the main structured domain of PrP, and are unstructured. Mutations in PRP1 are therefore unlikely to affect the structure of PrP^C^, and may exert their effects through misfolded PrP, other intermediate states, or altered protein-protein interactions. Similarly, the C-terminus of H2, also displays elevated conformational variability relative to other structural elements in the C-terminal domain [67-69]. It is markedly less stable than the remainder of H2 [70], and, depending on solution conditions, adopts an imperfect helical conformation as seen in a number of NMR structures [58, 71]. Its structural integrity is not mandatory for prion replication, as insertions within this region have no limiting effect on prion replication [72], although insertions affecting the charge interaction between the disease-associated residue H186 and K193 do inhibit prion propagation [72]. Alanine substitutions in this region of the protein are thus unlikely to significantly affect the structure of this region of PrP^C^ but may act through intermediate states, altered protein-protein interactions, or may affect the structure and stability of PrP in its pathologic conformation.

### Loss of prion propagation is more severe for mutations in H1, rather than H2 or H3

Propagation was weakest for PRP1 and PRP13. This was followed by PRP4 (S134A.R135A.M153A) and PRP5 (H139A.G141A.D146A), with mutations in H1 and neighbouring residues, reporting the next lowest indices of prion propagation (Fig. 1A & 3B). Both PRP4 and PRP5 showed a severe reduction in RML prion propagation (Fig. 3A).

Mutants that exclusively targeted H2 and H3 (PRP12, PRP14, PRP15 and PRP16, Fig.1A & 4B), although severely reduced in their ability to propagate RML prions, permitted sustained prion propagation and reported spot numbers higher than those observed for mutants PRP1 and PRP13, as well as PRP4 and PRP5, which are within and adjacent to H1 (Fig. 3B). We were unable to assess the effect of mutagenesis in the central H1 residues of the protein, mutant PRP7, as mutations in this region destroyed the ICSM18 epitope, used for the SCA (Fig. 2A). Overall, mutations to the surface regions of the protein (PRP1-PRP16) resulted in a severe loss of prion propagation, regardless of the region mutated.

### Mutations to core hydrophobic residues in PrP^C^ lower propagation

Set 3 mutations were replacements in the core hydrophobic region of the prion protein, including those thought to have destabilising and unfolding properties: PRP17 (M204A), PRP18 (M204Q.M205Q.V209A.M212Q) and PRP19 (C178A.C213A), which removed the PrP disulfide bridge (Fig.1A & 5B) [49, 73, 74]. Each of these mutants demonstrated a significant loss in prion propagation potential but generally reported higher spot numbers than mutants of surface residues, PRP1-PRP16. Mutants PRP17 and PRP18, but not PRP19 (Fig.5A), yielded higher spot numbers for the earliest time point at which they were assayed (split 4) compared to subsequent splits, which is the opposite of typical propagation. One possibility is that the higher spot count at the earlier time point was due to residual inoculum, but this was unlikely, as residual RML prions were not detected in the non-reconstituted PK1-KD cells at the equivalent time point (Fig. 5A). Alternatively, the observed spot numbers for PRP17, PRP18 and PRP19 could result from the destabilising mutations contributing to aberrant protein aggregation as opposed to *bona fide* propagation. To test this hypothesis, the SCA data for RML-infected and non-infected cells was compared. Non-RML infected cells exhibited fewer spot numbers than RML infected cells, indicating that the greater spot numbers reflected prion propagation and not aberrant protein aggregation. PRP19 exhibited low level propagation, whereas PRP17 and PRP18 appeared to exhibit only acute prion infection - a concept described by Vorberg *et al* [75], where a response to prion challenge is achieved in the infected cells, demonstrable by higher spot numbers relative to non-infected cells, but propagation is not maintained in consecutive splits within the SCA. This could be due to prion propagation being slower in these mutants than cell division such that prions are diluted out.

## Discussion

In this study, we employed site-directed alanine mutagenesis of surface residues within the structured domain (121-230) of the mouse prion protein to identify a ‘patch’ required for prion propagation. All mutations within the structured domain, irrespective of secondary structure, severely reduced prion propagation. The reduction was exacerbated for mutants targeting conformationally variable regions (PRP1 and PRP13) and those neighbouring or within H1 (PRP4 and PRP5). These sites may represent essential surfaces on PrP^C^ for template-assisted conversion following prion infection or may disrupt the prion structure itself as discussed below. Our data provide evidence in support of the three previously proposed regions that are important for successful infection and propagation of prions: the glycine rich region [41]; H1 and the neighbouring residues [28]; and the core residues in the protein that provide stability [59, 76]. These results corroborate previous studies that have identified the C-terminus of H2 as a key region in the conversion of PrP^C^ to its more β-sheet disease-associated form [48, 77]. Pathogenic mutations associated with inherited prion disease are also found in this region: T183A, H187R, T188R/K/A, E196K, F198S, E200K, and D202N [78-80].

### Conformationally variable regions within the structured domain of the cellular prion protein are the major determinants for the efficacy of prion propagation

Complete abrogation of propagation was observed for mutant 1 and PRP13 (Figs. 3 & 4), which lie on the same face of the protein’s globular domain. When raw spot numbers were analysed, there was a marginal but progressive drop with successive splits; split 6 values were below those observed for non-reconstituted PK1-KD cells. This suggested that the face of the globular domain in moPrP that bears these regions is an important site for prion propagation. Point mutations G122A and L124A, severely affect the ability of cells to take up and propagate prions [41]. Mutation G125A, also present in PRP1, was previously shown to offer partial resistance to prion infection, but not to the same extent as G122A and L124A [41]. Glycine to alanine mutations at positions G123A and G125A (PRP1) are predicted to disrupt the hairpin loop in the interlobe interface of the prion fibril and thus destabilize the amyloid fold (Fig. 3C) [11]. It remains to be investigated whether these sites are also required for contact between the native and disease-associated forms, native protein unfolding, association with propagation-facilitating factors, or are important residues for the stability of the disease-associated PrP assemblies. This glycine-rich region has been proposed to have lipid bilayer properties that may allow the protein to engage in transmembrane helix-helix interactions at this site [81]. In support of a conformational change in this loop region, it has been shown that the anti-prion monoclonal antibody 1C5, whose epitope spans residues 119-130, can recognise abnormal prion protein more effectively than the normal prion protein [82].

Structural effects of the PRP13 mutations on RML prion fibril are less obvious (Fig. 4C), suggesting that these mutations affect the misfolding pathway of PrP rather than the stability of the prion fibril. In PRP13, the region affected is the C-terminus of H2 (residues 187-194), which despite its high sequence conservation displays a high degree of conformational variability. NMR data indicate that the dominant α-helical structure of this region of the protein is unstable and interchanges with other lower populated structures [20, 70, 83]. Indeed, Salamat *et al*. have shown that peptide insertions here have no limiting effect on PrP^C^ conversion, and concluded that structural integrity within this region was not mandatory for successful prion propagation [72]. However, peptide insertion at K193, but not V188 or T190 in the loop region between H2 and H3, inhibited propagation [72]. Our data showed a strong restriction on prion propagation in mutant PRP13 (V188A.T191A.T192A). This region is important in terms of prion disease, as the disease-associated mutation of the neighbouring residue H187 to arginine has a dramatic effect on PrP^C^ folding and stability, resulting in a molecule that displays a markedly increased propensity to oligomerise [77]. Additionally, there are four consecutive threonine residues in this region (residues 190-193) that are found almost exclusively in a strand and/or loop conformation (irrespective of the identity of the flanking amino acids) [84].

Residues at the C-terminus of H2, clustered around H187 and T188, display high conformational flexibility, which enables a transition from α-helical to β-sheet or random coil states. This region contains many so-called ‘α/β discordant’ residues, proposed to be associated with amyloid formation [39]. Dynamic simulation of pathogenic mutations showed specific perturbation of loop H2-H3 for all mutations tested [85]. Our data from PRP13 highlighted native residues V188, T191, and T192 within the C-terminus of H2 as having a key role in prion propagation (Fig. 4A). Whether viewed from peptide insertion data, dynamic simulation or SCA, it was evident that prion propagation can be modulated by the C-terminus of H2 and the H2-H3 loop.

As conformationally variable regions are not likely to have a significant effect on the structure or stability of PrP^C^, or in the case of residues 122-126, lie outside the folded domain of PrP. These findings suggest that they may be exerting their effects on the propagative potential downstream of PrP^C^, for example, through a more rapid formation or increased stability of disease-associated PrP.

### H1 as a potential site for prion conversion

The data also strongly suggest that residues in and around H1 have an important role in prion propagation, as both PRP4 (S134A.R135A.M153A) and PRP5 (H139A.G141A.D146A) exhibited a marked reduction in prion propagation (Fig. 3A). These mutations replace polar residues on the inside of the N-terminal lobe of the prion fibril, which may disrupt the hydrogen bonding network in this region. However, the conformationally variable H1 region has previously been shown to play a crucial role in prion propagation in cell-free conversion studies; aspartate residues and salt-bridges, within H1, were thought to protect PrP^C^ from conversion through increased helix stability of the native state [33]. Residues neighbouring and within H1 (143-153) have been proposed as interaction sites where disease-associated PrP interacts with PrP^C^ [50]. Additionally, altering the charge orientation in this region impacts the success of prion formation [28]. H1 is a highly immunogenic region of the protein, and many anti-prion antibodies such as ICSM18 have their epitopes within H1; ICSM18 recognises PrP^C^ and disease-associated forms of PrP upon denaturation [86]. The data presented here suggest that H1 may play an important role in prion propagation, but does not act alone, as low levels of propagation were still detected upon H1 mutation (Fig. 3). Moreover, residues 143 and 146 within H1 have been reported as critical for propagation [87], with residues N-terminal to this region found to exhibit little control over propagation [88]. Previous work, probing conformational changes to the protein surface via chemical modifiers, showed H1 and H1-proximal residues to be more prone to modification and therefore more accessible in PrP^C^, relative to disease-associated PrP [89-91]. Molecular modelling of prevalent human prion protein mutations also indicated H1 as a region with the greatest degree of conformational change [92]. Movement of the H1 segment of PrP away from the H2-H3 core during prion conversion has been suggested previously [93, 94]. Our findings support the general idea that mutations, within and around H1, have a negative impact on propagation. However, they contradict the notion that regions N-terminal to it lack a strong influence on propagation, as a profound loss of propagation is observed for mutations in PRP1, in which residues 123-125 were mutated (Fig. 3A) in accordance with previous findings [41].

### Mutations in helices 2 and 3 do not abrogate, but severely inhibit propagation

Loss of prion propagation was less severe for mutations in H2 and H3 than for those in H1 and S1/S2 regions (Figs. 1 and 3). This was surprising since a large number of pathogenic mutations are clustered in H3 (22). This study has suggested that mutations at conformationally variable regions exert stronger control over prion propagation than those targeting surface regions on structured helices or beta-strands. Of the helical regions, H1 had a stronger influence over propagation outcome than H2 or H3, or even S1/S2 (Fig. 3 and 4).

### Do changes in protein stability affect prion propagation?

The alanine replacements targeted all secondary structure elements within PrP (Fig.1). Of these mutants, substitution of core hydrophobic residues least affected prion propagation (Fig. 5A). Mutations to the hydrophobic core destabilize native PrP^C^ and accelerate misfolding [36, 48]. Following the reasoning that lower native state stability correlates with altered folding intermediates and increased misfolding propensity [37], mutations to core residues should increase prion propagation. However, PRP12, PRP14, PRP15, which target specifically core residues, did not enhance prion replication (Fig. 4A). Similarly, oxidation of methionine residues perturbs the structural core of PrP^C^ [76]. However, SCA results showed that substituting methionine with alanine at M204, or altering residues M204.M205.V209.M212 by glutamine replacement, as in PRP18, resulted in considerably reduced prion propagation (Fig. 5A). Their prion propagation efficiency decreased with subsequent splits, and may be examples of mutant PrPs being incapable of sustaining infection [75]. Similarly, destabilization of PrP^C^ through removal of the disulfide bridge (PRP19) markedly reduced prion propagation (Fig. 5A), suggesting that the stability of PrP^C^ is not a major factor in prion replication efficacy.

### Implications for PrP^C^ structural changes on prion propagation

Taken together, the data presented indicate that structural mutations strongly decrease prion propagation, which may suggest that the same residues are crucial for the structure of the infectious prion or intermediate states involved in their production. Our mutational analysis also highlights key surface regions that have a strong influence on propagation efficiency, as evidenced by mutations in residues 123-125, 188-192 or 134-153 (Figs.1, 3 & 4).

When mapped onto PDB entry 2L39, sites 123-125 and 188-192 lie on the same face of the protein with region 134-153 on the opposite face (Fig. 6). It has been proposed that during prion conversion, there may be a conformational change in the S1-H1-S2 region away from the H2-H3 core of the prion protein [89, 90]. The S1-H2-S2 segment may be unfolded preferentially, whereas the H2-H3 segment remains marginally stable [95]. A shift away from the H2-H3 core may bring H1 and its neighbouring residues in proximity to the loop regions of PrP^C^. Such a mechanism would complement the SCA data that highlights residues 123-125, the C-terminus of H2 and H2-H3 loop, and H1, as the regions that most severely compromise prion propagation when mutated. These faces of the protein may act as important sites either as scaffolds for, or have direct involvement in, the interaction of PrP^C^ with prions resulting in successful propagation. Alternatively, residues 123-125 lie in a core region of recently determined prion structures [9-12], at the intersection between the two main lobes of the prion, where the introduction of bulkier side chain amino acids will likely destabilise the prion structure. The amino acid changes for residues 134-153 and 188-192 are more surface accessible and may therefore be more likely to be involved in the direct interaction of PrP^C^ with prions.

**Figure 6.**
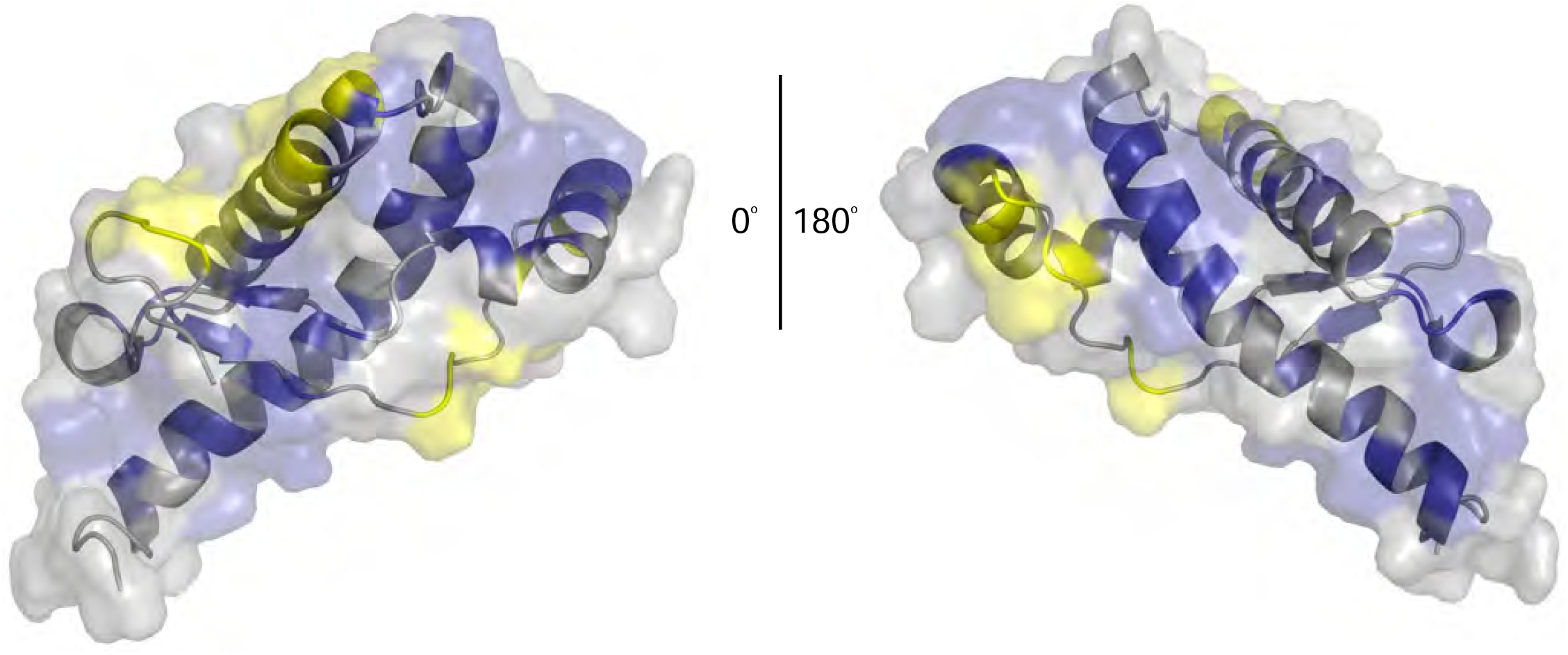
Surface residues of the prion protein that contribute to propagation efficiency. Regions that most severely compromised propagation following targeted mutagenesis are highlighted in yellow on PDB entry 2L39 rotated 180°C at the vertical axis. Other targeted regions are coloured in blue.

### Experimental methods

#### Construction of plasmid DNAs

pBluescript SK+ plasmid vector containing the full-length open reading frame (ORF) of mouse prion protein was used for mutagenesis. Mutations to alanine (codon GCC) were made using the Stratagene QuikChange® site-directed mutagenesis kit (Agilent Technologies, Santa Clara, USA). Mutants were sequence-verified and the ORFs inserted into the retroviral vector pLNCX2 (Takara Clontech, Mountain View, CA, USA).

#### Retroviral expression of alanine-mutants

10μg of pLNCX2 DNA containing the mutated ORFs were used to transfect phoenix ecotropic packaging cells (ATCC, LGC Standards, Middlesex, UK) with FuGENE 6 transfection reagent (Roche) according to the manufacturer’s instructions.

Supernatants were collected 24hrs post-transfection, passed through a 0.45μm filter and applied to PK1-KD cells. Cells stably transduced with the pLNCX2 constructs were selected using 200μg per ml G418 (Life Technologies, Thermofisher Scientific). Suppression of endogenous prion protein expression was achieved through stable expression of pRetroSuper retroviral vector [96] containing a shRNA sequence targeting the 3’UTR of the endogenous prion protein [55, 56]; stable silencing was ensured by maintaining the cells under puromycin selection. Stably transduced PK1-KD cells were selected using 200μg/ml G418 and 4μg/ml puromycin. Drug resistant colonies were pooled to obtain bulk cultures.

#### Scrapie Cell Assay

Scrapie cell assay (SCA) was carried out as previously described by Kloehn *et al* [18]. Briefly, cells were plated at 18000 cells/ well of a 96-well plate and Rocky Mountain Laboratory (RML) prion infected mouse brain homogenate added at 6309 tissue culture infectious units (TCIU; equivalent to a 10^−4^ dilution of 10% RML prion infected mouse brain homogenate) the following day. Cells were grown for three weeks with biweekly 1:8 splits. At splits 4, 5 and 6, cell suspensions equivalent to 25,000 cells were plated onto activated ELISPOT plates, lysed, digested with Proteinase K and probed with the mouse anti-PrP ICSM18 antibody (1:6,000 of 1mg/ml; D-Gen Ltd, UK) and goat anti-mouse anti-IgG1-AP secondary antibody (1:10,000 of 1mg/ml; Southern Biotech) to identify the number of PrP^Sc^ positive cells.

Quantitation of cells positive for PrP^Sc^ was carried out using WellScan software (Imaging Associates, Oxfordshire, UK). Object size of the detection module was set to 9 and the threshold (detection sensitivity) adjusted until all visible spots were detected; shape factor was set to 0.4 to exclusively detect spherical objects. Optimum values for threshold were found to be 25 to 30; a value of 28 was used in this study.

#### Immunofluorescence analysis

Sterile 22mm glass coverslips were coated with 1mg/ml Poly-L-Lysine for 30 min, washed with 1xPBS in the hood and dried, prior to seeding cells at 20,000 cells per coverslip. Cells were grown at 37°C in 5% CO_2_ and fixed using 4% w/v paraformaldehyde (PFA) for 20 min at RT. Cells were washed 3 times with 1xPBS, permeabilised with 0.05% Triton X100 and then incubated with the mouse anti-PrP ICSM18 primary antibody (1:7,000 of 1mg/ml) for 1 hour at RT, prior to 3 further washes with 1xPBS. AlexaFluor 488-conjugated goat anti-mouse was subsequently added at 1:1,000 for 1h at RT. After three washes with 1xPBS, coverslips were mounted using ProLong® Gold Antifade Reagent (Life Technologies, Thermofisher Scientific) containing DAPI for nuclear staining, and left to dry. Coverslips were stored at 4°C prior to imaging.

#### Image acquisition

Fluorescence images were acquired using a Zeiss LSM510 META laser scanning confocal microscope with a 40×/1.30 oil DIC objective, equipped with ConfoCor 3 detection module. Scanning was carried out at room temperature using λex = 405 nm (nuclear stain: DAPI) or 488 nm (mouse anti-PrP antibodies ICSM18 or ICSM35 and goat-anti-mouse Alexa-488 conjugated secondary antibody).

#### Western Blotting

Cell lysates were prepared from frozen cell pellets enriching for membrane associated proteins. Cells were resuspended in 10mM phosphate buffer (P5244, SigmaAldrich), incubated on ice for 5 mins (minutes) followed by centrifugation at 15,000g for 15 mins. The pellet was resuspended in D-PBS, digested with benzonase (1-2μl, 25KU equivalent to >250units/μl) for 15 mins at room temperature and boiled for 10 mins after addition of an equal volume of 2x sample buffer (125mM TrisHCl pH6.8, 20% v/v glycerol, 4% w/v SDS, 4% v/v 2-mercaptoethanol, 8mM 4-(2-aminoethyl)-benzene sulfonyl fluoride and 0.02 % w/v bromophenol blue). After centrifugation at 15000g for 1 min, the supernatant was removed and protein concentration determined by the Bradford Assay.

15-25μg of protein was fractionated on 12% BisTris NUPAGE gel (NP0341, ThermoScientific), transferred to PVDF and probed using anti-PrP ICSM35 antibody (0.2μg/ml, D-Gen LTD, London, UK). Antigen-antibody complexes were identified using goat anti-mouse AP (1:10,000 dilution of A2179, Sigma-Aldrich) and CDP-*Star*™ Substrate (T2146, ThermoScientific).

## ACKNOWLEDGEMENTS

We thank Ray Young and Richard Newton for graphics. This work was funded by the UKRI Medical Research Council.

## REFERENCES

[1] Collinge J. Prion diseases of humans and animals: their causes and molecular basis. Annual Review of Neuroscience. 2001;24:519–50.

[2] Gajdusek DC. Transmissible and non-transmissible amyloidoses: autocatalytic post-translational conversion of host precursor proteins to beta-pleated sheet configurations. J Neuroimmunol. 1988;20:95–110.

[3] Come JH, Fraser PE, Lansbury PTJ. A kinetic model for amyloid formation in the prion diseases: importance of seeding. Proc Natl Acad Sci USA. 1993;90:5959–63.

[4] Prusiner SB. Prions. Proc Natl Acad Sci USA. 1998;95:13363–83.

[5] Collinge J, Clarke A. A general model of prion strains and their pathogenicity. Science. 2007;318:930–6.

[6] Terry C, Wenborn A, Gros N, Sells J, Joiner S, Hosszu LL, et al. Ex vivo mammalian prions are formed of paired double helical prion protein fibrils. Open Biol. 2016;6:160035.

[7] Collinge J. Mammalian prions and their wider relevance in neurodegenerative diseases. Nature. 2016;539:217–26.

[8] Wenborn A, Terry C, Gros N, Joiner S, D’Castro L, Panico S, et al. A novel and rapid method for obtaining high titre intact prion strains from mammalian brain. Sci Rep. 2015;5:10062.

[9] Hoyt F, Alam P, Artikis E, Schwartz CL, Hughson AG, Race B, et al. 2022.

[10] Manka SW, Wenborn A, Betts J, Joiner S, Saibil HR, Collinge J, et al. A structural basis for prion strain diversity. Nat Chem Biol. 2023 doi:10.1038/s41589-022-01229-7.

[11] Manka SW, Zhang W, Wenborn A, Betts J, Joiner S, Saibil HR, et al. 2.7□Å cryo-EM structure of ex vivo RML prion fibrils. Nature Communications. 2022;13:4004.

[12] Kraus A, Hoyt F, Schwartz CL, Hansen B, Artikis E, Hughson AG, et al. High-resolution structure and strain comparison of infectious mammalian prions. Mol Cell. 2021.

[13] Meyer RK, McKinley MP, Bowman K, Braunfeld MB, Barry RA, Prusiner SB. Separation and properties of cellular and scrapie prion proteins. Proc Natl Acad Sci USA. 1986;83:2310–4.

[14] Safar J, Wille H, Itri V, Groth D, Serban H, Torchia M, et al. Eight prion strains have PrP^Sc^ molecules with different conformations. Nature Medicine. 1998;4:1157–65.

[15] Leske H, Hornemann S, Herrmann US, Zhu C, Dametto P, Li B, et al. Protease resistance of infectious prions is suppressed by removal of a single atom in the cellular prion protein. PLoS ONE. 2017;12:e0170503.

[16] Cronier S, Gros N, Tattum MH, Jackson GS, Clarke AR, Collinge J, et al. Detection and characterization of proteinase K-sensitive disease-related prion protein with thermolysin. Biochem J. 2008;416:297–305.

[17] Khalili-Shirazi A, Kaisar M, Mallinson G, s J, Bhelt D, Fraser C, et al. Beta-PrP form of human prion protein stimulates production of monoclonal antibodies to epitope 91-110 that recognise native PrP(Sc). Biochim Biophys Acta. 2007;1774: 1438–50.

[18] Klohn P, Stoltze L, Flechsig E, Enari M, Weissmann C. A quantitative, highly sensitive cell-based infectivity assay for mouse scrapie prions. Proc Natl Acad Sci USA. 2003;100:11666–71.

[19] Riek R, Hornemann S, Wider G, Billeter M, Glockshuber R, Wuthrich K. NMR structure of the mouse prion protein domain PrP (121-231). Nature. 1996;382:180–2.

[20] Hosszu LL, Jackson GS, C T, S J, Batchelor M, Bhelt D, et al. The residue 129 polymorphism in human prion protein does not confer susceptibility to CJD by altering the structure or global stability of PrP^C^. J Biol Chem. 2004;279:28515–21.

[21] Petsch B, Muller-Schiffmann A, Lehle A, Zirdum E, Prikulis I, Kuhn F, et al. Biological effects and use of PrPSc- and PrP-specific antibodies generated by immunizing with purified full length native mouse prions. J Virol. 2011; 85: 4538–4546.

[22] Doolan KM, Colby DW. Conformation-dependent epitopes recognized by prion protein antibodies probed using mutational scanning and deep sequencing. J Mol Biol. 2015;427:328–40.

[23] Lloyd S, Mead S, Collinge J. Genetics of Prion Disease. Topics in Current Chemistry. 2011;305:1–22.

[24] Ilc G, Giachin G, Jaremko M, Jaremko L, Benetti F, Plavec J, et al. NMR Structure of the Human Prion Protein with the Pathological Q212P Mutation Reveals Unique Structural Features. PLoS ONE. 2010;5:e11715.

[25] Bett C, Fernandez-Borges N, Kurt TD, Lucero M, Nilsson KP, Castilla J, et al. Structure of the beta2-alpha2 loop and interspecies prion transmission. FASEB J. 2012; 26: 2868–2876.

[26] Kurt TD, Bett C, Fernandez-Borges N, Joshi-Barr S, Hornemann S, Rulicke T, et al. Prion Transmission Prevented by Modifying the beta2-alpha2 Loop Structure of Host PrPC. J Neurosci. 2014;34:1022–7.

[27] Kurt TD, Jiang L, Fernandez-Borges N, Bett C, Liu J, Yang T, et al. Human prion protein sequence elements impede cross-species chronic wasting disease transmission. J Clin Invest. 2015;125(4):1485–1496.

[28] Norstrom EM, Mastrianni JA. The charge structure of helix 1 in the prion protein regulates conversion to pathogenic PrPSc. J Virol. 2006;80:8521–9.

[29] Ostapchenko VG, Makarava N, Savtchenko R, Baskakov IV. The Polybasic N-Terminal Region of the Prion Protein Controls the Physical Properties of Both the Cellular and Fibrillar Forms of PrP. J Mol Biol. 2008;383:1210–1224.

[30] Biljan I, Giachin G, Ilc G, Zhukov I, Plavec J, Legname G. Structural basis for the protective effect of the human prion protein carrying the dominant-negative E219K polymorphism. Biochem J. 2012;446:243–51.

[31] Guo J, Ren H, Ning L, Liu H, Yao X. Exploring structural and thermodynamic stabilities of human prion protein pathogenic mutants D202N, E211Q and Q217R. J Struct Biol. 2012; 178:225–232.

[32] Turnbaugh JA, Unterberger U, Saa P, Massignan T, Fluharty BR, Bowman FP, et al. The N-terminal, polybasic region of PrP(C) dictates the efficiency of prion propagation by binding to PrP(Sc). J Neurosci. 2012;32:8817–30.

[33] Speare JO, Rush TS, III, Bloom ME, Caughey B. The role of helix 1 aspartates and salt bridges in the stability and conversion of prion protein. J Biol Chem. 2003; 278:12522–12529.

[34] Kong Q, Mills JL, Kundu B, Li X, Qing L, Surewicz K, et al. Thermodynamic Stabilization of the Folded Domain of Prion Protein Inhibits Prion Infection in Vivo. Cell Rep. 2013 ; 4:248–254.

[35] Benetti F, Biarnes X, Attanasio F, Giachin G, Rizzarelli E, Legname G. Structural determinants in prion protein folding and stability. J Mol Biol. 2014; 426:3796–3810.

[36] Ning L, Guo J, Jin N, Liu H, Yao X. The role of Cys179-Cys214 disulfide bond in the stability and folding of prion protein: insights from molecular dynamics simulations. J Mol Model. 2014;20:2106.

[37] Benetti F, Legname G. New insights into structural determinants of prion protein folding and stability. Prion. 2015;9:119–124.

[38] Fernandez-Escamilla AM, Rousseau F, Schymkowitz J, Serrano L. Prediction of sequence-dependent and mutational effects on the aggregation of peptides and proteins. Nat Biotechnol. 2004; 22:1302–1306.

[39] Paivio A, Nordling E, Kallberg Y, Thyberg J, Johansson J. Stabilization of discordant helices in amyloid fibril-forming proteins. Protein Sci. 2004;13:1251–9.

[40] Yamaguchi K, Matsumoto T, Kuwata K. Critical region for amyloid fibril formation of mouse prion protein: unusual amyloidogenic properties of the helix 2 peptide. Biochemistry. 2008;47:13242–51.

[41] Harrison CF, Lawson VA, Coleman BM, Kim YS, Masters CL, Cappai R, et al. Conservation of a glycine-rich region in the prion protein is required for uptake of prion infectivity. J Biol Chem. 2010;285:20213–23.

[42] Giachin G, Mai PT, Tran TH, Salzano G, Benetti F, Migliorati V, et al. The non-octarepeat copper binding site of the prion protein is a key regulator of prion conversion. Sci Rep. 2015;5:15253.

[43] Beringue V, Mallinson G, Kaisar M, Tayebi M, Sattar Z, Jackson GS, et al. Regional heterogeneity of cellular prion protein isoforms in the mouse brain. Brain. 2003;126:2065–73.

[44] Ma J. The role of cofactors in prion propagation and infectivity. PLoS Pathog. 2012;8:e1002589.

[45] Yamaguchi KI, Kamatari YO, Fukuoka M, Miyaji R, Kuwata K. Nearly Reversible Conformational Change of Amyloid Fibrils as Revealed by pH-Jump Experiments. Biochemistry. 2013; 47:13242–13251.

[46] Tang Q, Fenton AW. Whole-protein alanine-scanning mutagenesis of allostery: A large percentage of a protein can contribute to mechanism. Hum Mutat. 2017;38: 1132–43.

[47] Fersht AR, Daggett V. Protein folding and unfolding at atomic resolution. Cell. 2002;108:573–82.

[48] Singh J, Udgaonkar JB. The Pathogenic Mutation T182A Converts the Prion Protein into a Molten Globule-like Conformation Whose Misfolding to Oligomers but Not to Fibrils Is Drastically Accelerated. Biochemistry. 2016;55:459–69.

[49] Hart T, Hosszu LL, Trevitt CR, Jackson GS, Waltho JP, Collinge J, et al. Folding kinetics of the human prion protein probed by temperature jump. Proc Natl Acad Sci USA. 2009;106:5651–6.

[50] Abalos GC, Cruite JT, Bellon A, Hemmers S, Akagi J, Mastrianni JA, et al. Identifying key components of the PrPC-PrPSc replicative interface. J Biol chem. 2008; 283:34021–34028.

[51] Hara H, Okemoto-Nakamura Y, Shinkai-Ouchi F, Hanada K, Yamakawa Y, Hagiwara K. Mouse prion protein (PrP) segment 100 to 104 regulates conversion of PrP(C) to PrP(Sc) in prion-infected neuroblastoma cells. J Virol. 2012;86:5626–36.

[52] Daskalov A, Gantner M, Walti MA, Schmidlin T, Chi CN, Wasmer C, et al. Contribution of Specific Residues of the beta-Solenoid Fold to HET-s Prion Function, Amyloid Structure and Stability. PLoS Pathog. 2014;10:e1004158.

[53] Ferguson N, Becker J, Tidow H, Tremmel S, Sharpe TD, Krause G, et al. General structural motifs of amyloid protofilaments. Proc Natl Acad Sci USA. 2006; 103:16248–16253.

[54] Williams AD, Shivaprasad S, Wetzel R. Alanine scanning mutagenesis of Abeta(1-40) amyloid fibril stability. J Mol Biol. 2006;357:1283–94.

[55] Bhamra S, Arora P, Manka SW, Schmidt C, Brown C, Rayner MLD, et al. Prion Propagation is Dependent on Key Amino Acids in Charge Cluster 2 within the Prion Protein. J Mol Biol. 2022;435:167925.

[56] Goold R, Rabbanian S, Sutton L, Andre R, Arora P, Moonga J, et al. Rapid cell-surface prion protein conversion revealed using a novel cell system. Nat Commun. 2011;2:281.

[57] Rogers M, Yehiely F, Scott M, Prusiner SB. Conversion of truncated and elongated prion proteins into the scrapie isoform in cultured cells. Proc Natl Acad Sci USA. 1993;90:3182–6.

[58] Damberger FF, Christen B, Perez DR, Hornemann S, Wuthrich K. Cellular prion protein conformation and function. Proc Natl Acad Sci USA. 2011;108:17308–13.

[59] Ning L, Wang Q, Zheng Y, Liu H, Yao X. Effects of the A117V mutation on the folding and aggregation of palindromic sequences (PrP113-120) in prion: insights from replica exchange molecular dynamics simulations. Mol Biosyst. 2014; 11:647–655.

[60] Halfmann R, Alberti S, Krishnan R, Lyle N, O’Donnell CW, King OD, et al. Opposing effects of glutamine and asparagine govern prion formation by intrinsically disordered proteins. Mol Cell. 2011;43:72–84.

[61] Caughey B, Raymond GJ. The scrapie-associated form of PrP is made from a cell surface precursor that is both protease- and phospholipase-sensitive. J Biol Chem. 1991;266:18217–23.

[62] Neuendorf E, Weber A, Saalmuller A, Schatzl H, Reifenberg K, Pfaff E, et al. Glycosylation deficiency at either one of the two glycan attachment sites of cellular prion protein preserves susceptibility to BSE and scrapie infections. J Biol chem. 2004;279:53306–16.

[63] Goold R, McKinnon C, Rabbanian S, Collinge J, Schiavo G, Tabrizi S. Alternative fates of newly formed PrPSc upon prion conversion on the plasma membrane. J Cell Sci. 2013; 126:3552–3562.

[64] Marijanovic Z, Caputo A, Campana V, Zurzolo C. Identification of an intracellular site of prion conversion. PLoS Pathog. 2009;5:e1000426.

[65] Enari M, Flechsig E, Weissmann C. Scrapie prion protein accumulation by scrapie-infected neuroblastoma cells abrogated by exposure to a prion protein antibody. Proc Nat Acad Sci USA. 2001;98:9295–9.

[66] Marbiah MM, Harvey A, West BT, Louzolo A, Banerjee P, Alden J, et al. Identification of a gene regulatory network associated with prion replication. EMBO J. 2014:e201387150.

[67] O’sullivan DB, Jones CE, Abdelraheim SR, Brazier MW, Toms H, Brown DR, et al. Dynamics of a truncated prion protein, PrP(113-231), from (15)N NMR relaxation: Order parameters calculated and slow conformational fluctuations localized to a distinct region. Protein Sci. 2008;18:410–23.

[68] Hosszu LLP, Conners R, Sangar D, Batchelor M, Sawyer EB, Fisher S, et al. Structural effects of the highly protective V127 polymorphism on human prion protein. Commun Biol. 2020;3:402.

[69] Dima RI, Thirumalai D. Probing the instabilities in the dynamics of helical fragments from mouse PrPC. Proc Natl Acad Sci USA. 2004;101:15335–40.

[70] Hosszu LLP, Baxter NJ, Jackson GS, Power A, a C, Waltho JP, et al. Structural mobility of the human prion protein probed by backbone hydrogen exchange. Nature Structural Biology. 1999;6:740–3.

[71] Calzolai L, Zahn R. Influence of pH on NMR structure and stability of the human prion protein globular domain. J Biol Chem. 2003;278:35592–6.

[72] Salamat K, Moudjou M, Chapuis J, Herzog L, Jaumain E, BÃ©ringue V, et al. Integrity of Helix 2-Helix 3 Domain of the PrP Protein Is Not Mandatory for Prion Replication*. J Biol Chem. 2012;287:18953–64.

[73] Maiti NR, Surewicz WK. The role of disulfide bridge in the folding and stability of the recombinant human prion protein. J Biol Chem. 2001;276:2427–31.

[74] Hirschberger T, Stork M, Schropp B, Winklhofer KF, Tatzelt J, Tavan P. Structural instability of the prion protein upon M205S/R mutations revealed by molecular dynamics simulations. Biophys J. 2006;90:3908–18.

[75] Vorberg I, Raines A, Priola SA. Acute formation of protease-resistant prion protein does not always lead to persistent scrapie infection in vitro. J Biol Chem. 2004; 279:29219–29225.

[76] Younan ND, Nadal RC, Davies P, Brown DR, Viles JH. Methionine oxidation perturbs the structural core of the prion protein and suggests a generic misfolding pathway. J Biol Chem. 2012;287:28263–75.

[77] Hosszu LL, Tattum MH, Jones S, Trevitt CR, Wells MA, Waltho JP, et al. The H187R mutation of the human prion protein induces conversion of recombinant prion protein to the PrP(Sc)-like form. Biochemistry. 2010;49:8729–38.

[78] Kuwata K, Nishida N, Matsumoto T, Kamatari YO, Hosokawa-Muto J, Kodama K, et al. Hot spots in prion protein for pathogenic conversion. Proc Natl Acad Sci USA. 2007;104:11921–6.

[79] Beck JA, Poulter M, Campbell TA, Adamson G, Uphill JB, Guerreiro R, et al. PRNP allelic series from 19 years of prion protein gene sequencing at the MRC Prion Unit. Hum Mutat. 2010;31:E1551–E63.

[80] Colby DW, Prusiner SB. Prions. Cold Spring Harb Perspect Biol. 2011;3: a006833.

[81] Cymer F, Veerappan A, Schneider D. Transmembrane helix-helix interactions are modulated by the sequence context and by lipid bilayer properties. Biochim Biophys Acta. 2012;1818:963–73.

[82] Choi JK, Park SJ, Jun YC, Oh JM, Jeong BH, Lee HP, et al. Generation of Monoclonal Antibody Recognized by the GXXXG Motif (Glycine Zipper) of Prion Protein. Hybridoma (Larchmt). 2006;25:271–7.

[83] Liu H, Farr-Jones S, Ulyanov NB, Llinas M, Marqusee S, Groth D, et al. Solution structure of syrian hamster prion protein (90-231). Biochemistry. 1999;38:5362–77.

[84] Dima RI, Thirumalai D. Proteins associated with diseases show enhanced sequence correlation between charged residues. Bioinformatics. 2004; 20:2345–2354.

[85] Meli M, Gasset M, Colombo G. Dynamic Diagnosis of Familial Prion Diseases Supports the beta2-alpha2 Loop as a Universal Interference Target. PLoS ONE. 2011;6:e19093.

[86] Khalili-Shirazi A, Kaisar M, Mallinson G, Jones S, Bhelt D, Fraser C, et al. Beta-PrP form of human prion protein stimulates production of monoclonal antibodies to epitope 91-110 that recognise native PrPSc. Biochim Biophys Acta. 2007;1774: 1438–50.

[87] Speare JO, Rush TS, III, Bloom ME, Caughey B. The role of helix 1 aspartates and salt bridges in the stability and conversion of prion protein. J Biol Chem. 2003; 278:12522–12529.

[88] Taguchi Y, Mistica AM, Kitamoto T, Schatzl HM. Critical significance of the region between Helix 1 and 2 for efficient dominant-negative inhibition by conversion-incompetent prion protein. PLoS Pathog. 2013;9:e1003466.

[89] Gong B, Ramos A, Vazquez-Fernandez E, Silva CJ, Alonso J, Liu Z, et al. Probing structural differences between PrPC and PrPSc by surface nitration and acetylation: evidence of conformational change in the C-terminus. Biochemistry. 2011; 50:4963–4972.

[90] Serpa JJ, Makepeace KA, Borchers TH, Wishart DS, Petrotchenko EV, Borchers CH. Using isotopically-coded hydrogen peroxide as a surface modification reagent for the structural characterization of prion protein aggregates. J Proteomics. 2013; 81:31–42.

[91] Serpa JJ, Makepeace KA, Borchers TH, Wishart DS, Petrotchenko EV, Borchers CH. Using isotopically-coded hydrogen peroxide as a surface modification reagent for the structural characterization of prion protein aggregates. J Proteomics. 2014;100:160–6.

[92] Behmard E, Abdolmaleki P, Asadabadi EB, Jahandideh S. Prevalent mutations of human prion protein: a molecular modeling and molecular dynamics study. J Biomol Struct Dyn. 2011;29:379–89.

[93] Cobb NJ, Sonnichsen FD, McHaourab H, Surewicz WK. Molecular architecture of human prion protein amyloid: a parallel, in-register beta-structure. Proc Natl Acad Sci U S A. 2007;104:18946–51.

[94] Lu BY, Chang JY. A 3-disulfide mutant of mouse prion protein expression, oxidative folding, reductive unfolding, conformational stability, aggregation and isomerization. Arch Biochem Biophys. 2007; 460:75–84.

[95] Honda RP, Yamaguchi KI, Kuwata K. Acid-induced Molten Globule State of a Prion Protein: Crucial Role of Strand 1-Helix 1-Strand 2 Segment. J Biol chem. 2014; 289:30355–30363.

[96] Berns K, Hijmans EM, Mullenders J, Brummelkamp TR, Velds A, Heimerikx M, et al. A large-scale RNAi screen in human cells identifies new components of the p53 pathway. Nature. 2004;428:431–7.

